# CRISPR-Cas9-mediated depletion of *O*-GlcNAc hydrolase and transferase for functional dissection of *O*-GlcNAcylation in human cells

**DOI:** 10.1101/2020.08.19.258079

**Authors:** Andrii Gorelik, Andrew T. Ferenbach

**Affiliations:** Department of Chemistry, Molecular Sciences Research Hub, White City Campus, Wood Lane, Imperial College London, London, W12 0BZ, UK; The Francis Crick Institute, 1 Midland Road, London, NW1 1AT, UK; Centre for Gene Regulation and Expression, School of Life Sciences, University of Dundee, Dow street, Dundee, DD1 5EH, UK

## Abstract

*O*-GlcNAcylation is an abundant post-translational modification (PTM) on serine and threonine residues of nuclear and cytoplasmic proteins. Although this PTM has been reported on thousands of proteins, *O*-GlcNAc transferase (OGT) and hydrolase (OGA) are the only two enzymes that perform the respective addition and removal of *O*-GlcNAc on protein substrates. To examine the consequences of deregulated *O*-GlcNAcylation, the *O*-GlcNAc field has mostly relied on the use of RNA interference to knockdown OGT/OGA and inhibitors to block their activities in cells. Here, we describe the first complete CRISPR-Cas9 knockouts of OGA and a knockdown of OGT (with a maximal decrease in expression of over 80%) in two human cell lines. Notably, constitutive depletion of one *O*-GlcNAc cycling enzyme not only led to a respective increase or decrease in total *O*-GlcNAcylation levels but also resulted in diminished expression of the opposing enzyme, as a compensatory mechanism, observed in previous short-term pharmacological studies. The OGA knockout system presents a convenient platform to dissect OGA mutations and was used to further characterise the single Ser405 *O*-GlcNAc site of human OGA using the *S*-GlcNAc genetic recoding approach, helping to identify an *S*-GlcNAc-specific antibody which was previously thought to primarily detect *O*-GlcNAc.

## Introduction

*O*-linked β-*N*-acetylglucosamine modification (*O*-GlcNAcylation) is an important and ubiquitous post-translational modification (PTM) mediated by the *O*-GlcNAc transferase (OGT) on serines/threonines of thousands of proteins residing in the cytoplasmic and nuclear compartments of metazoan cells^1–3^. The reverse reaction, *O*-GlcNAc hydrolysis, catalysed by the *O*-GlcNAc hydrolase (OGA), makes it a highly dynamic modification. OGT has been shown to be essential in mice^4^ and fruit flies^5^, but whilst OGA is dispensable in flies^6^, OGA knockout leads to developmental abnormalities and death of neonates in mice^7^. Disruption of *O*-GlcNAc signalling is linked to diseases including diabetes, neurodegeneration and cancer^8^.

OGT and OGA expression is reciprocally controlled in response to changes of intracellular *O*-GlcNAcylation levels in order to maintain *O*-GlcNAc homeostasis and to prevent aberrant signalling. For instance, it has been shown that OGT inhibition leads to a markedly decreased expression of OGA^9–15^. Since OGA expression is also modulated by the total levels of *O*-GlcNAc modification, treatment of cells with an OGA inhibitor increases *O*-GlcNAc levels, concomitantly elevating the expression of OGA and decreasing OGT expression as a compensatory mechanism^15^.

At a post-translational level, OGT is phosphorylated on several residues, some of which promote its nuclear localisation and alter substrate preference^16^, while others modulate *O*-GlcNAc transferase activity^17,18^. OGT is *O*-GlcNAcylated with low stoichiometry (<5%) on multiple sites but the function of these sites is mostly unclear^19,20^. Interestingly, histone demethylase LSD2 possesses E3 ubiquitin ligase activity that targets OGT for proteasomal degradation^21^. LSD2 was shown to be downregulated in various cancer cell lines, explaining why OGT expression is frequently upregulated in many types of cancer^8,21^. Ubiquitylation sites on OGT remain to be identified. Much less is known about how OGA is regulated by PTMs. During apoptosis OGA undergoes cleavage by caspase-3 at Asp413^22^, though the biological consequences of this processing are unknown. OGA is also modified with *O*-GlcNAc at Ser405 which reduces its stability in mouse embryonic stem cells (mESCs)^19,23^.

The roles of PTMs on OGA and OGT remain to be investigated in detail. However, overexpression of mutant constructs to pinpoint the site-specific functions of these PTMs in cells already expressing wild type OGT or OGA would be unlikely to produce a clear result. CRISPR-Cas9 genome editing technology can be used to overcome this issue. Yet, upon CRISPR-Cas9 gene editing, the efficiency of homology directed repair (to obtain knock-in mutations) is much lower than the non-homologous end-joining (to obtain gene knockouts), complicating the generation of immortalised human cell lines with knock-in mutations. This is especially complicated in cancer cell lines that often have more than two copies of a gene of interest. Therefore, knockout models of OGA and OGT for reconstitution studies would provide a rapid way of testing how certain OGA/OGT mutations complement these knockouts.

Importantly, in rescue experiments, expression of tagged constructs in OGA/OGT knockouts would allow convenient visualisation by western blot and immunofluorescence using established commercial antibodies against tags such as FLAG, HA or GFP, to overcome the limited availability of high-quality anti-OGT/OGA antibodies (in addition to validation of the currently available OGT/OGA antibodies). Such tools would also be beneficial for expressing mutant proteins, including those linked to diseases^24^, and as controls for OGA/OGT inhibitors^25^. Additionally, OGA and OGT knockouts in human cells would provide an opportunity to investigate mutual regulation and effects on *O*-GlcNAc homeostasis.

Immortalised mouse embryonic fibroblasts (MEFs) derived from OGA knockout mice and a hydroxytamoxifen inducible OGT-knockdown MEF line have been reported^26^. However, due to the essential nature of OGT, the viability of the latter MEF cell line is compromised upon hydroxytamoxifen induction in long-term experiments (>72 h). The single report of using CRISPR-Cas9 in relation to OGA is a partial (~50%) knockdown of OGA in mantle cell lymphoma cell lines Jeko-1 and Granta-519^27^. CRISPR-Cas9 genome editing technology has not been utilised to obtain homozygous OGA knockouts or to stably knockdown OGT in human cells.

This report describes the first example of complete OGA knockouts and stable OGT knockdowns (maximal decrease in expression of over 80%) in HeLa and HEK293 human cell lines, generated by CRISPR-Cas9. The data presented here indicate that the reciprocal feedback mechanism between OGA and OGT seen with inhibitors and RNAi knockdowns, extends to constitutive abrogation or reduction in their respective expression levels, and hence the respective *O*-GlcNAc hydrolase and transferase activities. OGA-null HEK293 cells were used as a platform to probe the role of human OGA *O*-GlcNAcylation on Ser405 by overexpressing the OGA^S405C^ mutant with a non-hydrolysable *O*-GlcNAc mimic, *S*-GlcNAc, using the previously reported genetic recoding approach^23,28^. This convenient system allowed testing of the effects of hyper-*S*-GlcNAcylation on OGA localisation and caspase-3 cleavage. Additionally, an *S*-GlcNAc-reactive antibody was identified which was previously thought to primarily detect *O*-GlcNAcylated proteins.

## Results

### CRISPR-Cas9-mediated depletion of OGT

A strategy to knockout OGT in human cells using CRISPR-Cas9 genome editing was devised based on the sizes of the first exon. Since the open reading frame (ORF) region of the first exon of OGT is too small for efficient deletion, the second exon was chosen as a gRNA target (Fig. 1a).

**Figure 1.**
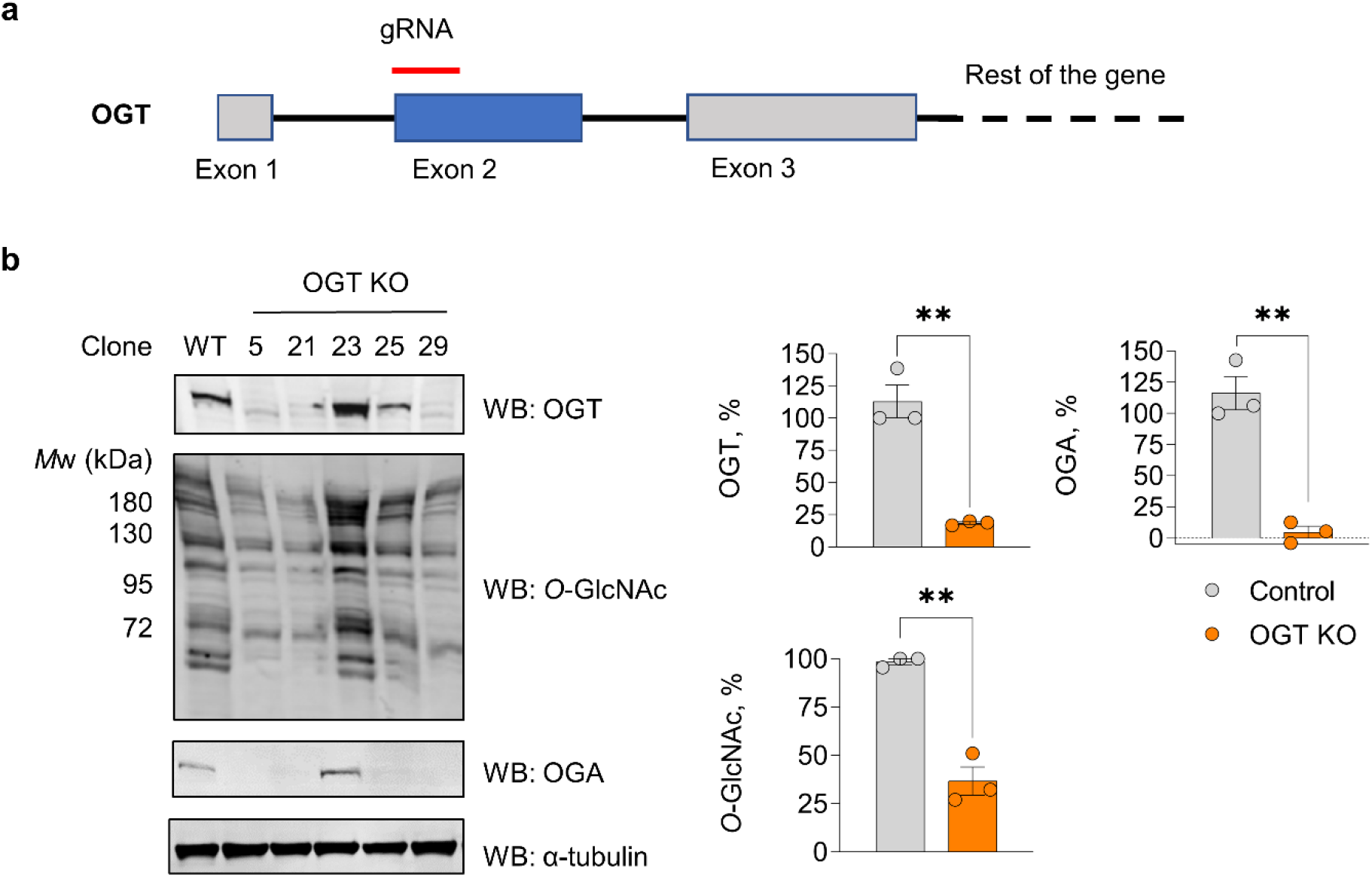
CRISPR-Cas9 knockdowns of OGT in HeLa cells. **a**, Strategy for the CRISPR-Cas9 knockout of OGT (OGT KO). **b**, OGT knockdown in HeLa cells results in decreased *O*-GlcNAcylation levels and abolished OGA expression, as detected by western blot using the corresponding antibodies. α-tubulin was used as a loading control. Error bars represent mean ± s.e.m. of three biological replicates, ** *P*<0.01, (*t*-test, two-tailed, unpaired).

CRISPR-Cas9 was then performed to knockout endogenous OGT in HeLa cells and several viable clones were obtained. Western blot analysis of cell lysates obtained from these clones indicated an efficient knockdown of OGT (>80%), alongside a surprising downward band-shift detected by an antibody raised against the first 300 amino acids of OGT (Fig. 1b). To investigate whether this band-shift could be caused by non-specific antibody binding, another anti-OGT antibody (against amino acids 213-462) was used. Similarly, the OGT band corresponding to a truncated protein was detected when probed with this different antibody (Supplementary Fig. 1).

*O*-GlcNAc levels in OGT-depleted HeLa cells were significantly reduced by almost three-fold (Fig. 1b). In accordance with the homeostatic regulation, OGA expression was abolished in OGT-depleted clones (Fig. 1b). In a rescue experiment, C-terminally HA-tagged OGT was overexpressed in OGT-depleted HeLa cells which led to an increase in total *O*-GlcNAcylation levels (Supplementary Fig. 2).

To investigate whether complete OGT knockout could be achieved in another cell line, HEK293 cells were subjected to CRISPR-Cas9 gene editing to delete OGT. In the case of the HEK293 cell line, only one clone was recovered which exhibited reduced OGT expression with molecular weight corresponding to full-length OGT (Supplementary Fig. 3a). Although the OGT depletion efficiency was less than that in HeLa cells, a substantial reduction in *O*-GlcNAc levels and abolished OGA expression were similarly observed (Supplementary Fig. 3a).

Interestingly, no obvious morphological or cell growth defects were observed in either HeLa or HEK293 OGT-depleted cells. Thus, OGT-depleted HeLa and HEK293 cell lines present a platform for investigating the mechanisms of *O*-GlcNAc homeostasis and for expressing various OGT mutants to test their activity, interactions and regulation.

### CRISPR-Cas9 knockout of OGA

Unlike OGT, the first exon of OGA is appropriate for gRNA targeting and was therefore targeted to knockout OGA in HeLa cells using CRISPR-Cas9 (Fig. 2a). Several viable clones were obtained with no detectable OGA expression as determined by western blot (Fig. 2b). As expected, *O*-GlcNAcylation levels in these cells were elevated (by approximately 2.5-fold) compared to wild type cells (Fig. 2b). OGT expression was also downregulated by about 50% in OGA knockout (OGA KO) clones (Fig. 2b), though not to such a great extent as the OGA expression in OGT knockout cells (Fig. 1b). Additionally, CRISPR-Cas9-mediated deletion of OGA was applied in HEK293 cells which yielded several clones (Supplementary Fig. 3b). Similar to HeLa OGA KO clones, these exhibited downregulated OGT expression and increased *O*-GlcNAcylation levels (Supplementary Fig. 3b). Neither HeLa nor HEK293 OGA KO cells exhibited any obvious morphological or cell growth defects. Therefore, OGA KO in HeLa and HEK293 cells present a convenient system for overexpression of OGA mutants.

**Figure 2.**
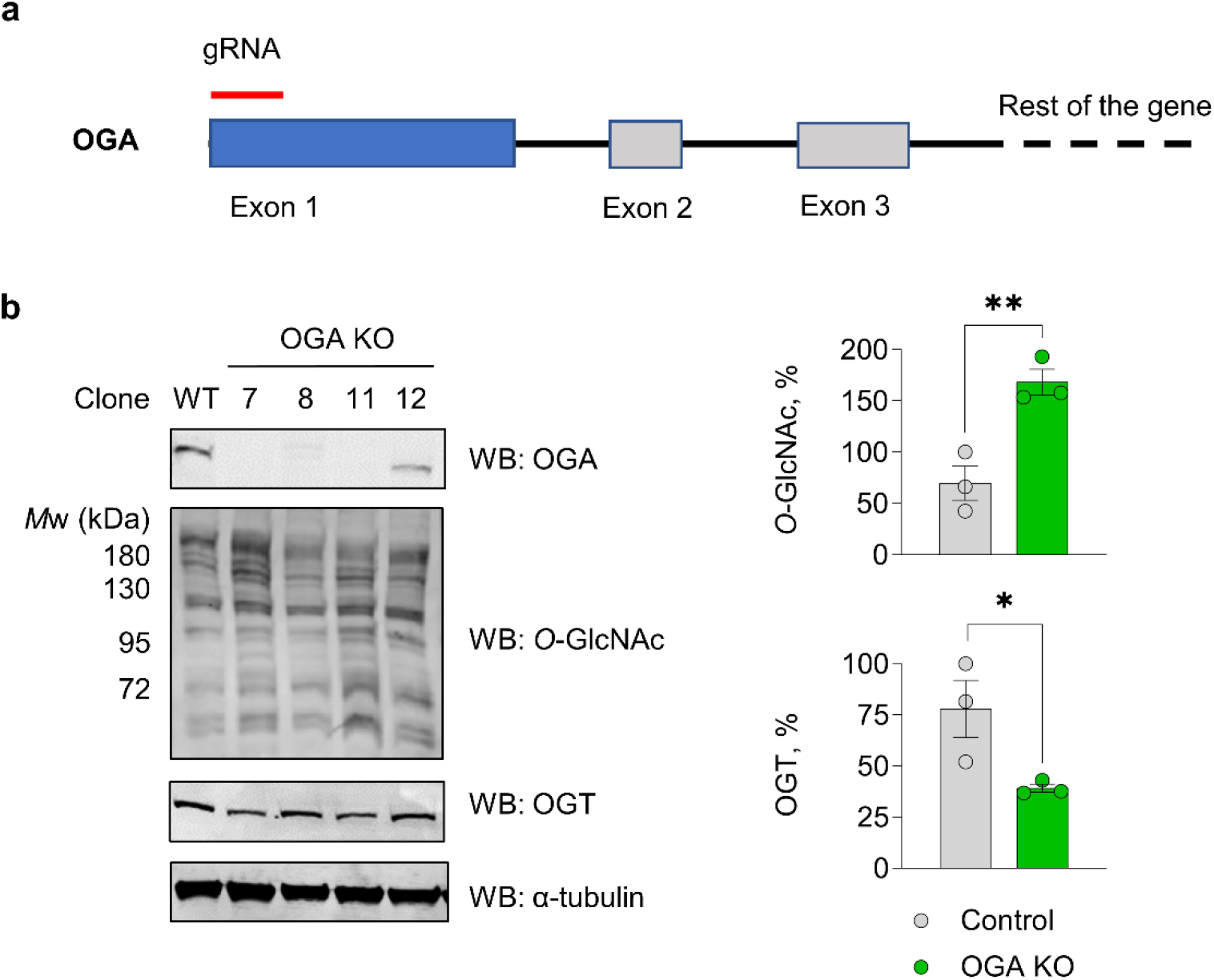
CRISPR-Cas9 knockouts of OGA in HeLa cells. **a**, Strategy for the CRISPR-Cas9 knockout of OGA (OGA KO). **b**, OGA knockout in HeLa cells results in increased *O*-GlcNAcylation levels and decreased OGT expression, as detected by western blot using the corresponding antibodies. OGA KO clones 7 and 11 were used for quantification. α-tubulin was used as a loading control. Error bars represent mean ± s.e.m. of three biological replicates, * *P*<0.05, ** *P*<0.01, (*t*-test, two-tailed, unpaired).

### Probing Ser405 *O*-GlcNAcylation of OGA in CRISPR-Cas9 OGA-null cells

Dissection of the roles of a single Ser405 *O*-GlcNAc site on OGA is complicated by the rapid off-cycling resulting in low stoichiometry^20^ and requires tools that site-specifically install a permanent *O*-GlcNAc modification without affecting OGT and OGA activities. Recently, genetic encoding of cysteines in place of natural serine/threonine *O*-GlcNAcylation sites allowed the incorporation of non-hydrolysable Cys-*S*-GlcNAc with high stoichiometry and site-specific precision^23^. Introduction of such an *S*-GlcNAc site in place of Ser405 of OGA in mESCs leads to hyper-GlcNAcylated OGA^23^. Application of this approach in combination with a cycloheximide chase assay led to the conclusion that increased *O*-GlcNAcylation of OGA reduces its stability^23^. However, it is possible that this modification of OGA plays other roles. Moreover, irreversible *S*-GlcNAcylation of S405C OGA in human cells remains to be demonstrated.

To further investigate the function of Ser405 *O*-GlcNAcylation site on human OGA, in a rescue experiment FLAG-tagged OGA^WT^ and OGA^S405C^ constructs were overexpressed in HEK293 OGA KO cells (clone 1, Supplementary Fig. 3b). Additionally, to prevent *O*-GlcNAc removal, cells overexpressing the OGA^WT^ construct were treated with GlcNAcstatin G, a validated nanomolar OGA inhibitor^29^, for 24 h prior to lysis. To quantify the stoichiometry of GlcNAc modification on OGA, galactosyltransferase GalT^Y289L^ labelling with GalNAz was performed in cell lysates followed by copper(I)-catalysed ligation to an alkyne-labelled PEG 5000 to allow separation of glycosylated and unmodified OGA by SDS-PAGE^20^. Quantification of GlcNAcylation stoichiometry revealed undetectable *O*-GlcNAcylation at a basal level, while the S405C mutation resulted in a GlcNAcylation stoichiometry of approximately 70%, similar to that induced by GlcNAcstatin G treatment on the wild type protein (Fig. 3a). Western blot analysis of total *O*-GlcNAcylation levels in the overexpressing cells revealed that the OGA^S405C^ mutant retains activity comparable to the wild type enzyme, despite being irreversibly hyper-*S*-GlcNAcylated in HEK293 OGA KO cells (Fig. 3b).

**Figure 3.**
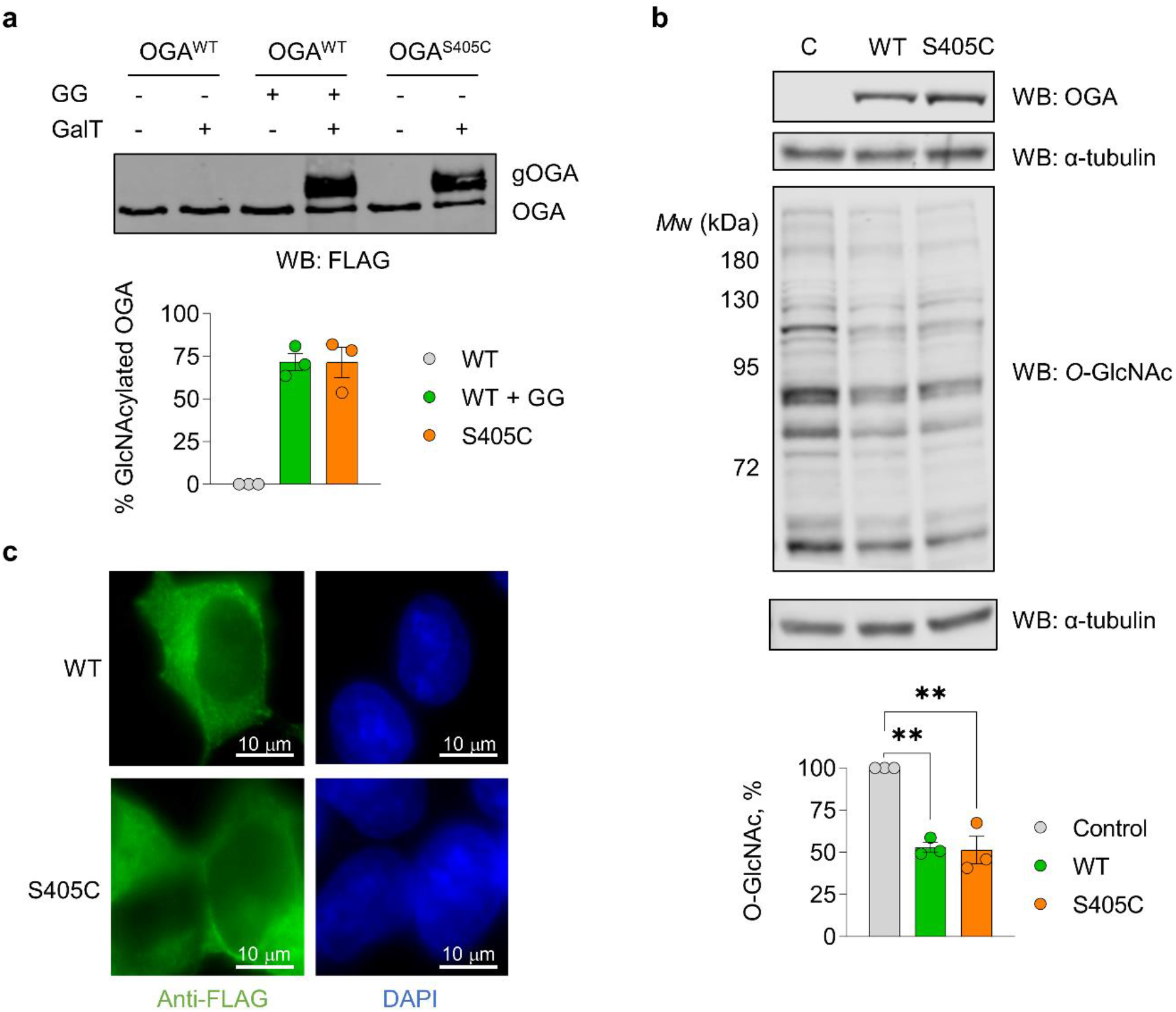
OGA knockout model to study hyper-GlcNAcylation of OGA. **a**, Stoichiometry of GlcNAcylation on WT and S405C FLAG-OGA, as detected by western blot. GG – GlcNAcstatin G treatment (1 μM, 24 h). gOGA – GlcNAcylated OGA. **b**, Overexpressed WT and S405C FLAG-OGA lead to decreased global *O*-GlcNAcylation in HEK293 OGA KO cells, as detected by western blot. C – untransfected control. α-tubulin was used as a loading control. **c**, Localisation of WT and S405C FLAG-OGA, as detected by immunofluorescence. DAPI staining was used to detect nuclei. In panels **a** and **b** error bars represent mean ± s.e.m. of three biological replicates, ** *P*<0.01, (*t*-test, two-tailed, unpaired).

Previously, Ala mutation of Ser399 *O*-GlcNAcylation site of OGT was shown to decrease its nuclear localisation^30^. To examine, whether the elevated GlcNAcylation stoichiometry could affect OGA localisation, immunofluorescence imaging was performed on cells overexpressing WT and S405C OGA with an N-terminal FLAG tag. However, when probed with an anti-FLAG antibody, no significant differences in localisation between WT and C405 OGA were observed with both mainly localised to the cytoplasm (Fig. 3c).

The FLAG tag provides an opportunity to pulldown OGA and analyse its GlcNAcylation status using the commercially available pan-specific anti-*O*-GlcNAc antibodies. To probe OGA GlcNAcylation, OGA was immunoprecipitated with an anti-FLAG antibody followed by a western blot analysis of the pulldowns with anti-OGA and pan-specific *O*-GlcNAc antibodies CTD110.6 and RL2, respectively. Strikingly, the S405C mutation resulted in the *S*-GlcNAcylation of OGA as detected by CTD110.6 (but not RL2), while the GlcNAcstatin G-induced *O*-GlcNAcylation was detected only by the RL2 antibody (Fig. 4a).

**Figure 4.**
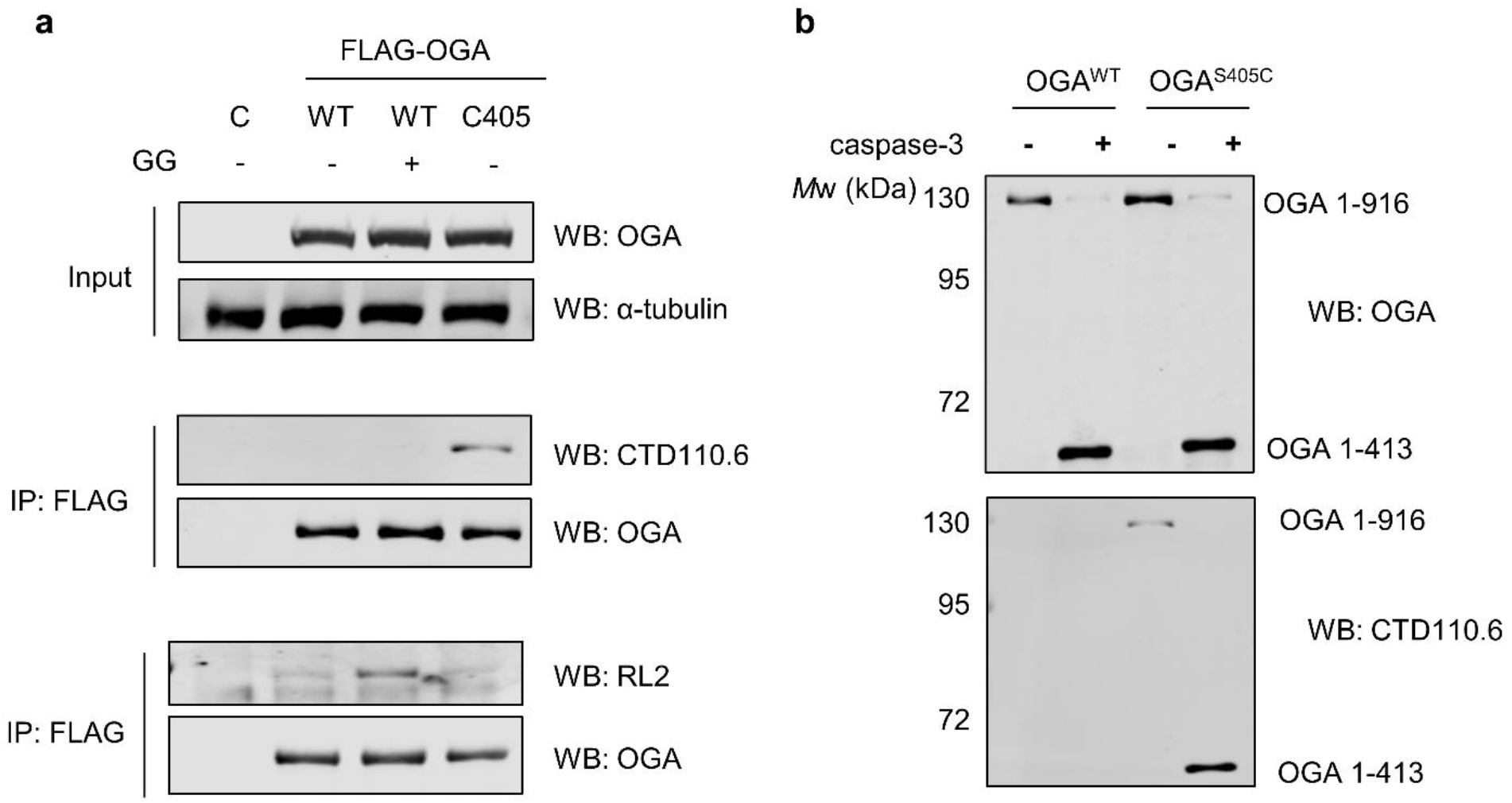
Detection of OGA *O*/*S*-GlcNAcylation and an *in vitro* caspase-3 cleavage assay on FLAG-OGA immunoprecipitated from cells. **a**, *O*-GlcNAcylation on WT OGA immunoprecipitated from HEK293 cells can be detected exclusively by the RL2 antibody while *S*-GlcNAcylation on S405C OGA can be detected only by the CTD110.6 antibody. GG – GlcNAcstatin G treatment (1 μM, 24 h). C – untransfected control. α-tubulin was used as a loading control. **b**, *In vitro* caspase-3 cleavage of immunoprecipitated OGA is not affected by increased GlcNAcylation stoichiometry, as detected by western blot. CTD110.6 antibody was used to detect *S*-GlcNAc.

Previously, proximal *O*-GlcNAc modification on caspase-8 has been shown to inhibit its auto-cleavage and subsequent activation^31^. To investigate whether high GlcNAcylation stoichiometry on Cys405 inhibits OGA^S405C^ cleavage by caspase-3 on Asp413, WT and S405C OGA immunoprecipitated from cells were subjected to cleavage by recombinant caspase-3. Despite the close proximity of the OGA^S405C^ *S*-GlcNAcylation site to the caspase-3 cleavage site, *in vitro* caspase-3 cleavage of both WT and hyper-GlcNAcylated S405C OGA proceeded to completion with the N-terminal OGA fragment being *S*-GlcNAcylated in the S405C mutant, as detected with the CTD110.6 antibody (Fig. 4b). These data further confirm the presence of *S*-GlcNAc modification in the N-terminal portion of OGA^S405C^ which encompasses Cys405.

## Discussion

Here, complete OGA knockouts were obtained in HeLa and HEK293 cells. Cell lines with over 80% depletion of OGT were obtained in HeLa cells and a less efficient knockdown in HEK293 cells, indicating that higher OGT activity may be more important for survival of some cell types than others. OGA knockout resulted in a substantial reduction of OGT expression, while disruption of OGT expression led to undetectable OGA levels. Previously it was shown that prolonged OGT inhibition leads to fewer productive OGA transcripts^32^, consistent with the CRISPR-Cas9 data presented in this manuscript which shows that OGA expression is abolished by OGT knockdown. Despite HeLa and HEK293 cell lines having at least three copies of either of the two *O*-GlcNAc cycling genes, CRISPR-Cas9 was shown to be highly efficient in removing them. Since OGT is essential, a small fraction of (truncated) OGT retained by HeLa cells likely enabled their survival.

OGA-null and OGT-depleted cell lines generated by CRISPR-Cas9 could help to define the mechanism of homeostatic regulation between OGA and OGT. It has been shown that intron retention between exons 4 and 5 (which encompasses a premature stop-codon) of the *OGT* transcript negatively regulates its expression in response to increased *O*-GlcNAc modification^32,33^. Similarly, transcriptional regulation of OGA expression via retained introns was reported as part of a feedback regulation loop^32^. *OGA* contains two introns between exons 10 and 12 split by exon 11 and these introns can undergo retention. However, contrary to OGT regulation, transient retention of these introns during splicing ultimately leads to a functional transcript and higher OGA expression in conditions of elevated *O*-GlcNAc modification, while exon 11 skipping and introduction of a premature stop-codon occurs at low *O*-GlcNAc levels leading to decreased OGA expression^32^. Importantly, the retained intron regions in OGA and OGT are conserved in vertebrates, implying a common mechanism^32^. Cell lines described here could help to identify the factor, or multiple factors (such as *O*-GlcNAc sensors), responsible for the regulation of OGA/OGT splicing and expression.

These cell lines provide an opportunity to investigate not only the mechanisms that regulate *O*-GlcNAc homeostasis but also systems for characterising OGA and OGT mutants. Additionally, they represent useful tools for the analysis of OGT/OGA inhibitors (i.e. in-cell screens of OGT inhibitors would identify promising hits that engage with a smaller amount of OGT relative to wild type cells, likely resulting in lower IC50 values required for inhibition)^24,25^. Here, this system was used to further characterise the hyper-GlcNAcylated OGA (S405C mutant) by *S*-GlcNAc mutagenesis^23,28^.

WT and S405C OGA exhibited similar activities in human cells confirming the previous findings in mouse embryonic stem cells^23^. *S*-GlcNAcylation stoichiometry on the S405C OGA in human cells was consistent with previous findings in mESCs^23^. Since the anti-FLAG antibody binding to the N-terminal FLAG tag does not interfere with PEGylation at residue 405 of OGA, this provided a clear western blot image and therefore a more accurate stoichiometry measurement. Basal *O*-GlcNAc stoichiometry was not detected in HEK293 cells unlike in our previous report in mESCs (15%)^23^, suggesting that the roles of this modification could be tissue-specific.

Introduction of a FLAG tag allowed the assessment of OGA localisation by immunofluorescence which is mainly cytoplasmic even when stoichiometric *S*-GlcNAcylation is introduced. FLAG tag also allowed a pulldown and detection of *O*/*S*-GlcNAc on OGA using RL2 and CTD110.6 antibodies. Interestingly, despite similar *O*- and *S*-GlcNAc stoichiometries, RL2 exclusively detected *O*-GlcNAc while CTD110.6 exclusively detected *S*-GlcNAc. Thus, antibodies have to be carefully chosen when detecting *O*-GlcNAc^28^. We have previously shown that CTD110.6 recognises *S*-GlcNAc on TAB1^S395C^ and CKII^S347C^ mutants^23^. CTD110.6 could be used to detect *S*-GlcNAc on individual (immunoprecipitated or purified) proteins, as well as total *S*-GlcNAc modification in cell lysates upon treatment with *Cp*OGA and PNGase F to remove *O*- and *N*-GlcNAc^34^. Very recently, using glycopeptide microarrays it was shown that CTD110.6 may preferentially bind *S*-GlcNAc in certain sequence contexts, supporting our findings with *O*- and *S*-GlcNAcylated OGA^35^.

Overall, CRISPR-Cas9 OGA knockouts and OGT-depleted cells present a new model to investigate the mechanisms underlying *O*-GlcNAc signalling and the factors that influence these.

## Acknowledgements

This work was funded by a Wellcome Trust 4-year PhD studentship (105310/Z/14/Z) to AG. We thank Dr Vladimir S. Borodkin for providing GlcNAcstatin G. We thank Dr Tony Ocasio (The Francis Crick Institute), Prof Ed Tate (Imperial College London), Dr Vladimir S. Borodkin (University of Dundee), Dr Olawale Raimi (University of Dundee) and Dr Sergio Galan Bartual (University of Dundee) for critical reading of the manuscript and useful suggestions.

## Author contributions

AG conceived the study and performed experiments, ATF performed cloning. AG wrote the manuscript with input from ATF.

## Conflict of interest

There are no conflicts of interest to declare

## Materials and Methods

### Generation of OGT knockdowns and OGA knockouts in Hela and HEK293 cells

Paired gRNA sites targeting the first exon of human OGA and the second exon of human OGT were chosen using the website http://www.sanger.ac.uk/htgt/wge/find_crisprs. Annealing oligonucleotides were designed with the appropriate overhangs for cloning into *BpiI* cut pX335 (Cas9 D10A) and pBABED puro U6. 3 μl of each annealing oligonucleotide were combined in a 100 μl volume and treated with 1 μl polynucleotide kinase (Fermentas) in T4 Ligase buffer at 37 °C for 20 min followed by 10 min incubation at 75 °C and finally placed in a heating block at 95 °C. The metal block was removed from the heat source and allowed to cool gradually to room temperature. 1 μl of a 1/30 dilution of this mixture was added to a ligation reaction containing 20 ng *Bpi*I cut, dephosphorylated destination vector and 1 μl DNA ligase (Fermentas) in T4 ligase buffer in a 20 μl final volume. After 25 min at room temperature, a 1 μl aliquot of the reaction was used to transform DH5-α competent cells. Inserts were confirmed by DNA sequencing. Primers used for knockout generation are listed in Table 1.

**Table 1.**
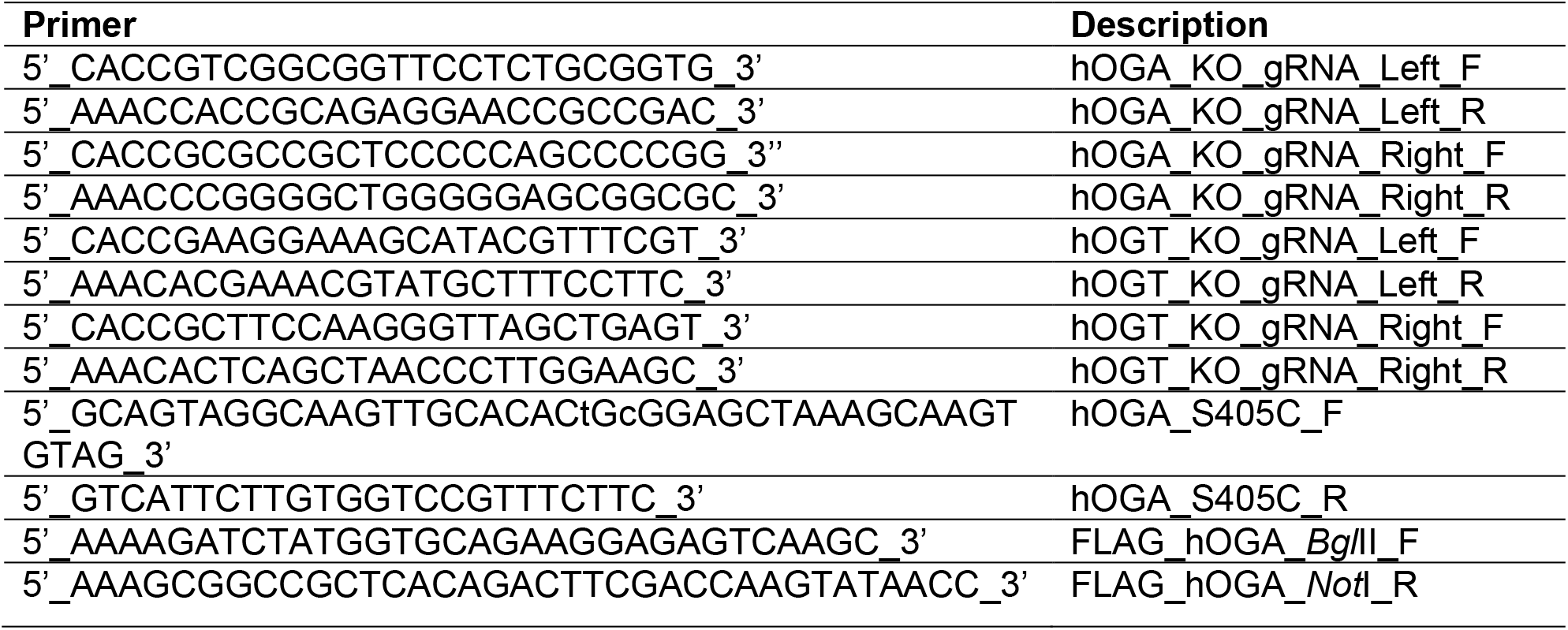
List of primers.

HEK293 or HeLa cells were co-transfected with pBABED and pX335 plasmids using Lipofectamine 2000 (Thermo Fisher Scientific) according to manufacturer’s instructions. 24 h post transfection, DMEM medium (supplemented with 10% FBS, 2 mM L-glutamine and 1% penicillin/streptomycin (100 U/ml and 100 μg/ml respectively)) was substituted with fresh medium containing 3.3 μg/ml puromycin. Cells that survived after 48 h of puromycin treatment were trypsinized, counted and single clones were plated on 96-well plates. Colonies were grown and passaged onto 6-well plates. Candidate clones were then tested by western blot with OGT- and OGA-specific antibodies to identify the knockout cell lines.

### Cloning

Gene coding for full-length OGT was cloned into pCMV-(C)-HA vector (obtained from DSTT, School of Life Sciences, Dundee, UK) for expression of the C-terminal HA tag. Gene coding for full-length human OGA was cloned into pCMV-FLAG vector (obtained from DSTT, School of Life Sciences, Dundee, UK) for expression of the N-terminal FLAG tag. Full-length human OGA containing the S405C mutation was produced by PCR, followed by restrictionless cloning using KOD polymerase from an existing WT construct. The products were gel extracted and digested with *BglII* and *NotI* restriction enzyme. They were then ligated into pCMV-FLAG cut with *BamHI* and *Not*I. The inserts were confirmed by DNA sequencing. Primers used for cloning and sequencing are listed in Table 1.

### Cell culture and transfection

HEK293 and HeLa cells were grown in DMEM medium supplemented with 10% FBS, 2 mM L-glutamine and 1% penicillin/streptomycin (100 U/ml and 100 μg/ml respectively) at 37 °C and 5% CO2. HEK293 cells were transfected with FLAG-tagged OGA using Lipofectamine 3000 reagent (Thermo Fisher Scientific) at a ratio 1:2 (μg:μl) according to manufacturer’s instructions. GlcNAcstatin G treatment was performed at a final concentration of 1 μM for 24 h, with DMSO as a vehicle control.

### Western blotting

Cells were scraped in lysis buffer (Cell Signaling Technology, #9803) supplemented with 1 mM PMSF, sonicated and lysates were spun down at 17,000 g for 10 min. Supernatants were transferred into fresh tubes and protein concentration was quantified using Bradford assay. Protein samples were prepared using LDS buffer containing 5% β-mercaptoethanol. Samples were run on a 10% SDS PAGE gel. Proteins were then transferred onto nitrocellulose membrane at 100 V for 1 h. Membranes were stained with Ponceau-S to check for successful transfer, washed with 0.2% TBS-Tween and blocked in 5% BSA in 0.2% TBS-Tween. Primary antibodies used: RL2 (ab2739. Abcam and MA1-072, Thermo Fisher Scientific, 1:1000), CTD110.6 (9875, CST, 1:1000) OGA (14711-1-AP, Proteintech 1:1000 – detects the first 350 amino acids of human OGA), OGT (H300, Santa-Cruz and ab96718, Abcam, 1:1000), α-tubulin (2125, CST, 1:1000). Fluorescence signal from secondary antibodies (LI-COR) was quantified using the Odyssey LI-COR system. Quantification was performed using Image Studio software (LI-COR) and statistical analysis was performed using Prism (GraphPad).

### Immunoprecipitation

Immunoprecipitation from HEK293 cell lysates was conducted using Dynabeads protein G magnetic beads (Invitrogen, Thermo Fisher Scientific) according to manufacturer’s protocol with a following modification: the lysates were first incubated with an anti-FLAG tag antibody at 4 °C overnight, followed by coupling of an antigen-bound antibody to the beads for 1.5 h at 4 °C.

### *O*- and S-GlcNAc PEGylation labelling using Gal-T^Y289L^

GalT^Y289L^ labelling and PEGylation were performed as previously described^23^.

### Immunofluorescence

OGA KO cells cultured on coverslips were transfected with FLAG-tagged WT OGA and S405C OGA plasmids. 24 h after transfection, the cells washed twice with Dulbecco’s Phosphate-Buffered Saline (D-PBS) and fixed with 4% formaldehyde for 20 min at room temperature. After two washes with D-PBS, cells were permeabilized with 0.1% Triton-X-100 in D-PBS for 10 min at room temperature. Cells were blocked with 5% BSA and incubated with a FLAG antibody (Sigma, mouse) at 1:500 dilution in 1% BSA, 0.1% Tween-20 in D-PBS at room temperature for 1 h. After three washes with 0.2% TBST, secondary antibody was added (anti-mouse Alexa Fluor ^®^ 488 1:1000) in 1% BSA (in D-PBS 0.1% Tween) and incubated for 1 h at room temperature. Coverslips were washed twice and DAPI (diluted 1:1000 v/v) was added during one of the washes for 1 min followed by three washes with 0.2% TBST. The coverslips were mounted onto glass slides using mounting medium (Dako) and imaged using DeltaVision widefield microscopy system.

### In vitro caspase-3 cleavage assay

Overexpressed FLAG-tagged OGA (WT or S405C mutant) was immunoprecipitated from HEK293 OGA knockout cells using an anti-FLAG antibody coupled to magnetic beads. Caspase-3 cleavage was performed on-beads using 0.25 units of recombinant caspase-3 (Enzo) for 4 h at 37 °C.

## Supplementary figures

**Supplementary figure 1.**
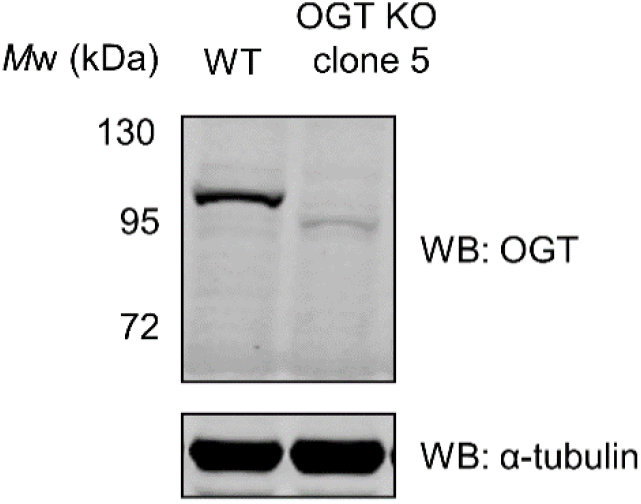
Detection of residual OGT in OGT KO HeLa cells. An antibody raised against residues 213-462 of OGT (Abcam) was used to detect truncated OGT by western blot in HeLa OGT KO clone 5 (Fig. 1b). α-tubulin was used as a loading control.

**Supplementary figure 2.**
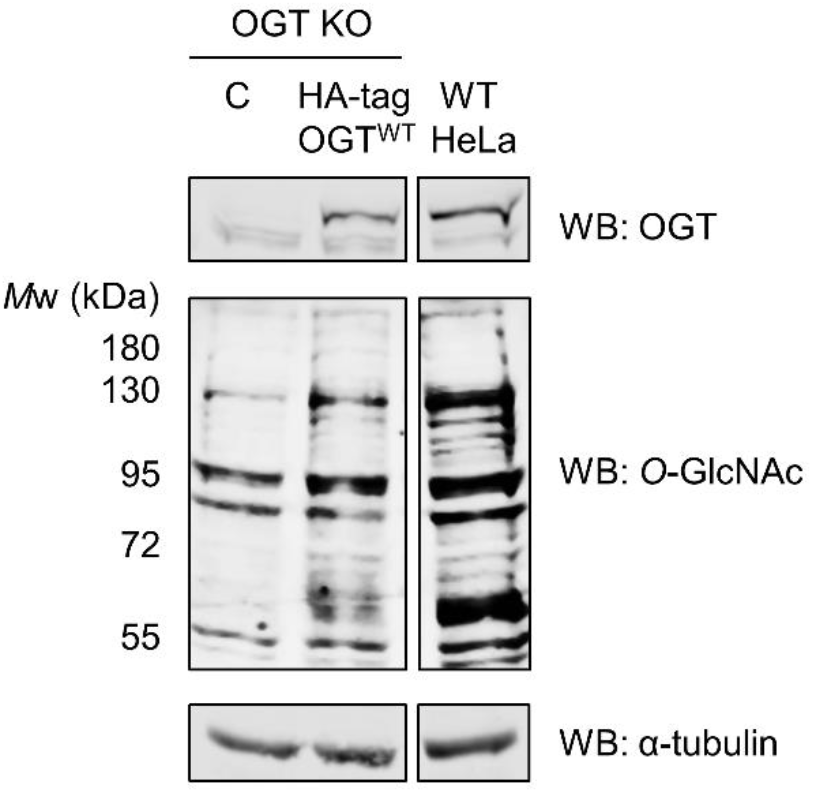
OGT overexpression in OGT KO HeLa cells elevates global *O*-GlcNAcylation levels. C-terminally HA-tagged OGT was overexpressed in OGT KO HeLa cells (clone 5 from Fig. 1b). OGT expression and *O*-GlcNAc levels were detected using the corresponding antibodies by western blot. α-tubulin was used as a loading control. C – untransfected control.

**Supplementary figure 3.**
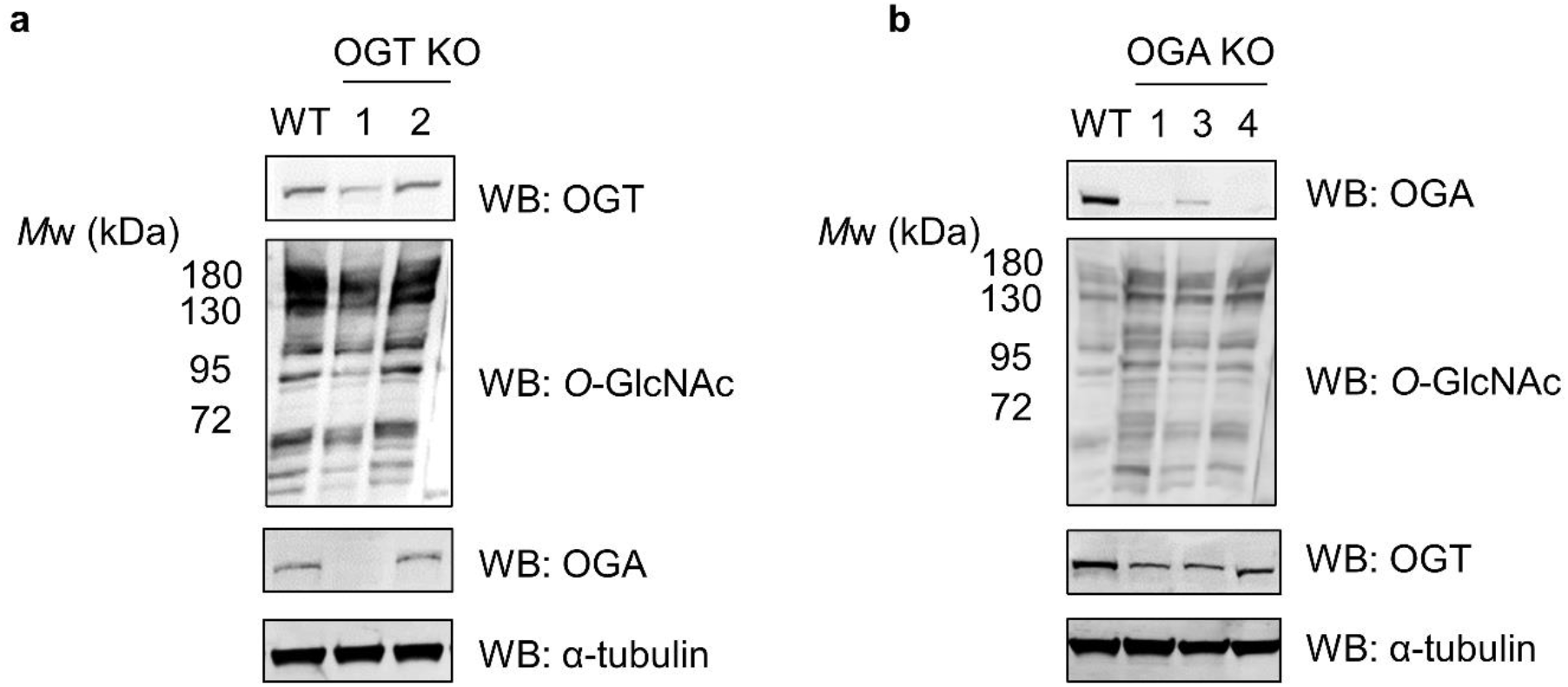
CRISPR-Cas9 knockouts of OGT and OGA in HEK293 cells. **a**, Incomplete OGT knockout (knockdown) was achieved in HEK293 cells resulting in decreased *O*-GlcNAcylation levels and abolished OGA expression, as detected by western blot. α-tubulin was used as a loading control. **b**, OGA knockout in HEK293 cells results in increased *O*-GlcNAcylation levels and decreased OGT expression, as detected by western blot. α-tubulin was used as a loading control.

## Notes

### Competing Interest Statement

The authors have declared no competing interest.

